# Molecular basis and evolutionary cost of a novel phenotype of macrolides/lincosamides resistance in *Staphylococcus haemolyticus*

**DOI:** 10.1101/2021.05.26.445780

**Authors:** Ruilin Xu, Tianqiang Song, Hong Du, Qiang Wang, Haifang Zhang

## Abstract

*Staphylococcus haemolyticus* (*S. haemolyticus*) is a coagulase-negative *Staphylococcus* and has become one of the primary pathogens of nosocomial infections. After a long period using the antibiotics against the infections caused by *S. haemolyticus, S. haemolyticus* developed several resistance phenotypes of macrolides and lincosamides. Here, we acquired four *S. haemolyticus* clinical isolates, of which three isolates demonstrated antibiotic resistance as reported previously, while one was resistant to both erythromycin and clindamycin without erythromycin induction. Such novel phenotype, known as constitutive Macrolides-Lincosamides-Streptogramins (MLS) resistance, was reported in other bacteria, but it was found for the first time in *S. haemolyticus*. To further investigate such behavior, we confirmed that the deletion of the methyltransferase *ErmC* upstream leader peptide was critical to cause constitutive MLS resistance based on the analysis of potential antibiotic resistance genes via whole-genome sequencing and experimental verification. In addition, we discovered that the continuous expression of *ErmC* may inhibit the growth of *S. haemolyticus*, which turned out to be the cost with no MLS pressure. Taken together, this study firstly reported the constitutive MLS resistance in *S. haemolyticus*, and provided new ideas for using macrolides and lincosamides in clinical treatment.

## 1. Introduction

*Staphylococcus* is a group of gram-positive bacteria that widely distributed in nature. According to the presence or absence of plasma coagulase, a critical pathogenic factor, *Staphylococcus* can be divided into coagulase-positive and -negative *Staphylococcus*. The former primarily refers to the virulent *S. aureus*, while the latter includes *S. epidermidis, S. haemolyticus, S. saprophytic*, et cetera. Coagulase-negative *Staphylococcus* (CoNS) has long been considered a harmless microbe [1]. Until the 1980s, with the widespread use of broad-spectrum antibiotics, the reports on various opportunistic infections caused by CoNS gradually increased [2, 3]. Moreover, due to the increasing usage of invasive diagnosis and artificial organ materials, CoNS has become one of the primary pathogens of nosocomial infections [4]. *S. haemolyticus*, which can produce hemolysins and enterotoxins [5], has strong pathogenicity for ICU patients, the elderly, newborn children, and patients with a defective immune system [6]. And it can easily cause sepsis, valve endocarditis, nosocomial acquired pneumonia, and urinary tract infection [7, 8]. It was reported that *S. haemolyticus* was the most frequent aetiological agent of staphylococcal infections together with *S. aureus* and *S. epidermidis* [9]. In the past 20 years, the clinical treatment of *S. haemolyticus* mainly included broad-spectrum penicillins, macrolides, lincosamides, first- and second-generation cephalosporins, aminoglycosides, quinolones, et cetera. With the increasing use of these antibiotics, the resistance of *S. haemolyticus* also increases year by year that causes more widespread nosocomial infections [10].

In the 1960s and 1970s, shortly after introducing erythromycin and clindamycin into clinical treatment, *S. haemolyticus* performed several corresponding phenotypes: resistance to erythromycin or clindamycin, respectively; resistance to erythromycin and clindamycin with erythromycin induction. The last one, known as inducible MLS phenotype, is the bacteria’s resistance to macrolides, lincosamides and streptogramins induced by macrolides [11]. For the MLS antibiotics, *S. haemolyticus* mainly has three resistance mechanisms. First, pumping antibiotics out of the cell by efflux pumps (a membrane transporter protein), that are mostly encoded by genes of ATP binding cassette (ABC) transport superfamily, including *Vga(A)*_*LC*_, *msrA* [12, 13]. Second, the inactivation of antibiotics, such as *mphC*, encodes a macrolide phosphotransferase that modifies antibiotics by phosphorylation, rendering them inactive [14]. Third, altering the antibiotics’ target, which prevents antibiotics from binding to targets. For instance, the ribosomal RNA methyltransferase *Erm* family, adding methyl modification to bacterial 23S rRNA renders the antibiotics unable to bind to 50S ribosomal large subunit and work effectively [15].

Recently, we explored four strains of *S. haemolyticus* separated by clinical infectious patients. Surprisingly, we identified a strain with constitutive MLS resistance that had not been reported previously. In this study, we investigated the molecular mechanism of such phenotype and confirmed by experiments. Such discovery not only broaden our understanding of bacteria evolution, but also provided suggestions in clinical treatment.

## 2. Materials and methods

### 2.1 Bacterial strains and media

*S. haemolyticus* strains A, B, C and D used in this study were initially isolated from patients with invasive clinical infections in the Second Hospital of Soochow University (Jiangsu, China). Standard *S. aureus* strain ATCC25923 were preserved in Soochow University. All original strains were grown in trypticase soy broth at 37 °C. Plasmid pBT2 (*Staphylococcus*-*Escherichia* shuttle vector) was used as a backbone [16]. Since pBT2 is a temperature-sensitive plasmid, it is easy to be lost when incubated at higher temperatures, all strains with pBT2 related plasmid were cultured at 30 °C.

### 2.2 Genome sequencing and assembly

We performed next-generation sequencing (Illumina, Hiseq2500, PE100) on these four isolates of *S. haemolyticus*. Raw reads were trimmed by Quorum, BBTools and Sickle. To achieve better assembly results, we integrate five tools to assemble clean reads: SuperReads from Masurca [17], BCALM [18], Tadpole from BBTools [19], SPAdes [20], and Megahit [21]. Assemblies of these tools were imported into Sequencher (Gene Codes, MI, USA), and the consensus sequences were manually verified and selected to form final assembly files.

### 2.3 Antibiotics resistance gene analysis

Prokka was used to annotate the assembled sequences [22]. The potential resistance genes were analyzed by Resistance Gene Identifier (RGI) developed by CARD [23]. According to the phenotypes, we found several genes related to erythromycin and clindamycin resistance, then compared them with their homologs in NCBI.

### 2.4 Plasmid construction and transformation

*ErmC* gene was amplified from the genomic DNA of *S. haemolyticus* strain C and inserted into the pBT2 backbone with digestion and ligation at restriction sites XbaI and EcoRI to form pBT2-*ErmC*, followed by sanger sequencing confirmation. The upstream leader peptide sequence (short for LP) of *ErmC* was obtained from *S. haemolyticus* strain PK-01 chromosome (NCBI Accession: CP035541 Region: 2,573,055-2,573,114) and synthesized by GenScript Biotech Co. The recombinant plasmid pBT2-LP-*ErmC* was constructed by In-Fusion Cloning. Briefly, homologous repeat sequences were designed at both ends of the recombinant plasmid pBT2-*ErmC* and the synthesized leader peptide. With the help of homologous recombinase (In-Fusion HD Cloning Kit, Takara), the two ends of the sequence with homologous arms were connected and verified by Sanger sequencing. Plasmids were transformed into *S. aureus* strain ATCC25923 by electrotransformation under the conditions of 2.0 kV, 25 μF and 100 Ω. All primers for plasmids construction (*ErmC*-XbaI-F, *ErmC*-EcoRI-R; In-Fusion vector-F/R, In-Fusion insert-F/R) used in this study were provided in Table S2.

### 2.5 Isolation of RNA and RT-PCR

The total RNA of wild-type *S. aureus* together with three experimental groups carried plasmid pBT2, pBT2-*ErmC* and pBT2-LP-*ErmC* (the latter two were cultured in medium with erythromycin), were extracted by the Trizol method (Invitrogen, Carlsbad, CA, USA). RNA purity and integrity were examined by RNA electrophoresis in 1.0% (w/v) agarose gel. One µg RNA was treated with DNase I prior to cDNA synthesis using a Revert-Aid First Strand cDNA Synthesis Kit (Thermo Scientific, MA, USA). RT-PCR was performed with specific primers of *ErmC* and *Staphylococcus* 16S rRNA fragment by electrophoresis in 2.0% (w/v) agarose gel, visualized with Tanon 3500. All RT-PCR primers (rt-*ErmC-*F/R, rt-*Staphylococcus* 16S-F/R) used in this study were provided in Table S3.

### 2.6 Antimicrobial susceptibility test

The turbidity of *S. aureus* transformed with recombinant plasmids was adjusted to 0.5 McFarland turbidity (1.5×10^8^ CFU/ml) and then coated onto Mueller-Hinton agar medium evenly for antimicrobial susceptibility test with sterilized cotton swabs. After 2 minutes, erythromycin susceptibility test disc (15 μg) and clindamycin susceptibility test disc (2 μg) were placed onto the middle of the medium, the distance between the disc and the edge of the Petrie dish was more than 15 mm, and the distance of the two discs was 15–26 mm apart, 30 °C cultured for 20–24 hours to check the results. The current CLSI M100 (Performance Standards for Antimicrobial Susceptibility Testing) determines whether the bacteria were resistant to the antibiotics.

### 2.7 Growth curve measurement

The *S. aureus* transformed with different plasmids were cultured at the same condition (30 °C, trypticase soy broth, chloramphenicol, 200 rpm/min). OD600 was measured every two hours in 20 h.

## 3. Results

### 3.1 Identifying a novel phenotype of macrolides/lincosamides resistance in *Staphylococcus haemolyticus*

In the Second Hospital of Soochow University, we isolated four strains of *S. haemolyticus* numbered A, B, C and D. During the routine drug sensitivity test to screen suitable antibiotics, strains A and B were resistant to clindamycin and sensitive to erythromycin; strain D was resistant to erythromycin and sensitive to clindamycin; strain C was resistant to both erythromycin and clindamycin, and was able to tolerate clindamycin directly without erythromycin induction, known as the constitutive MLS resistance. The phenotypes of strain A, B, and D were common in clinical practice, and their potential resistance mechanisms were elucidated entirely previously. However, the phenotype of strain C that we were never reported before.

The constitutive MLS resistance has been reported in *Staphylococcus aureus* [24, 25], *S. epidermidis* [26], *Streptococcus pyogenes* [27], *S. pneumoniae* and *Enterococcus faecalis* [28]. However, no similar phenotype was reported in *S. haemolyticus*. In the following work, we tried to figure out the reasons that led to such antibiotic resistance.

### 3.2 Genome assembly and antibiotic resistance gene analysis

The genomes of four *S. haemolyticus* clinical isolates were sequenced and assembled. The assembly results were listed in table 1, and all four assemblies demonstrated high quality and high coverage. Then the potential resistance genes were analyzed by RGI and showed in Table S1. Based on previous studies, several genes were identified to be related to erythromycin and clindamycin resistance: *Vga(A)*_*LC*_, *msrA, mphC* and *ErmC. S. haemolyticus* strains A and B contained *Vga(A)*_*LC*_; strain C contained *msrA, mphC* and *ErmC*; strain D contained *msrA* and *mphC* as shown in Fig 1. As expected, the antibiotic resistance genes existed in strains A, B, D were corresponding with their phenotypes. Due to the specific phenotype of strain C compared with others, we focused on exploring the cause of its formation. After comparing the sequences of the above resistance genes from strain C with their homologs in NCBI, we found a short leader peptide was missing from the upstream of *ErmC* (Fig 2). Then we looked into the reports about *ErmC* in the database (Table S2), and found that more of the *ErmC* gene had intact upstream leader peptide and the majority of them were usually located on plasmids. In addition, this type of leader peptide mutation identified in our study was the first to be found in *S. haemolyticus* and the first occur on chromosome. Further investigation showed the variation of leader peptide could render downstream *ErmC* gene to switch from inducible expression to constitutive expression, which resulted in constitutive MLS resistance (Fig 3) [26, 29, 30]. Based on this result, we hypothesized that such phenotype manifested by *S. haemolyticus* strain C was caused by the leader peptide deletion upstream of *ErmC*.

**Table 1.** Genome assembly results of the four *S. haemolyticus* strains.

| Strain | Reads Number | N50 | Contigs | Total Length | BUSCO Complete(%) |
| --- | --- | --- | --- | --- | --- |
| A | 12932836 | 97887 | 41 | 2572850 | 100.0 |
| B | 13832808 | 84894 | 47 | 2426878 | 100.0 |
| C | 8770750 | 66588 | 63 | 2652164 | 99.2 |
| D | 13103638 | 90527 | 52 | 2596861 | 100.0 |

**FIG 1.**
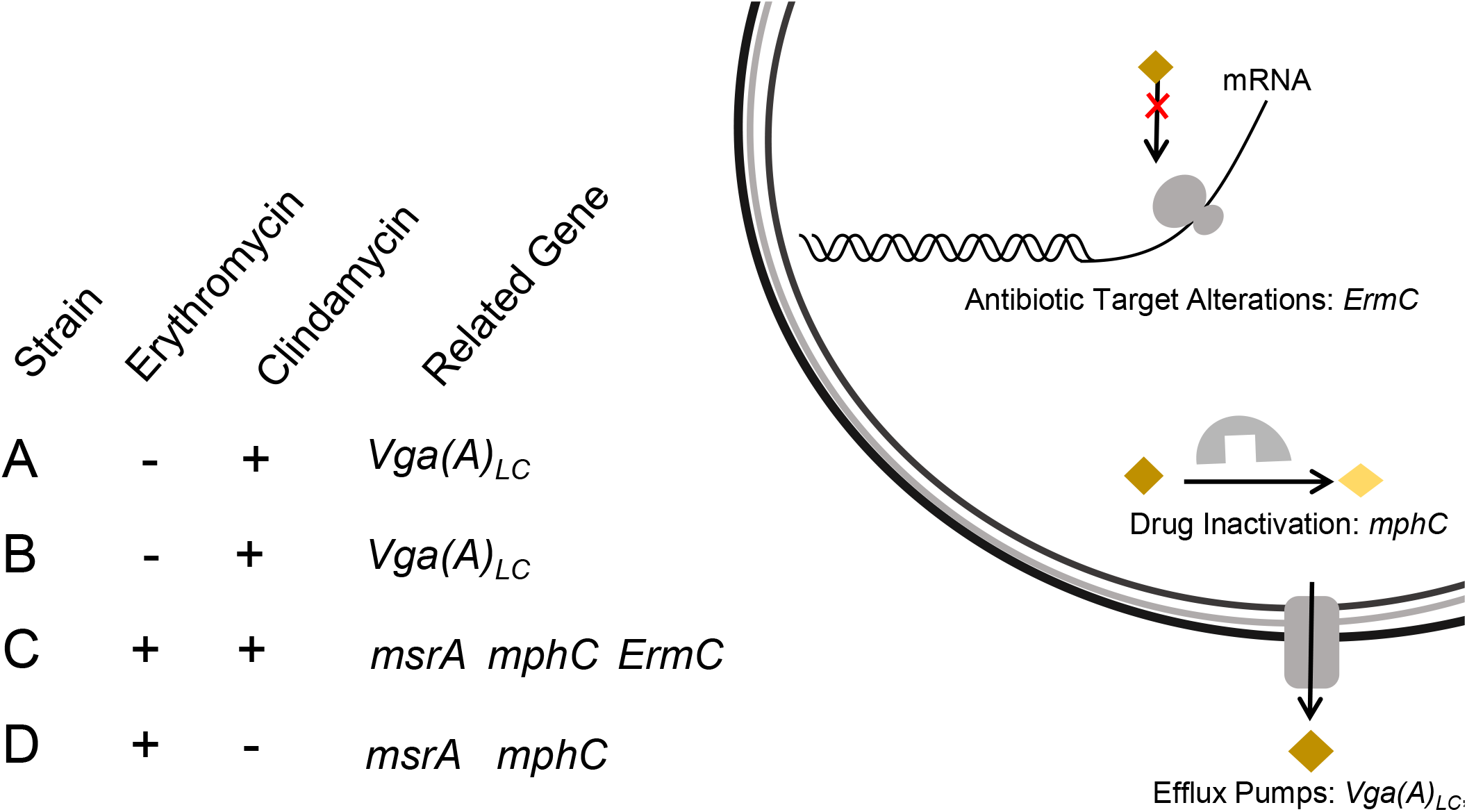
The resistance genes related to erythromycin and clindamycin detected from four *S. haemolyticus* strains ABCD. The plus or minus represented strains were resistant or sensitive to antibiotics. And the resistance mechanisms of these genes were shown on right.

**FIG 2.**
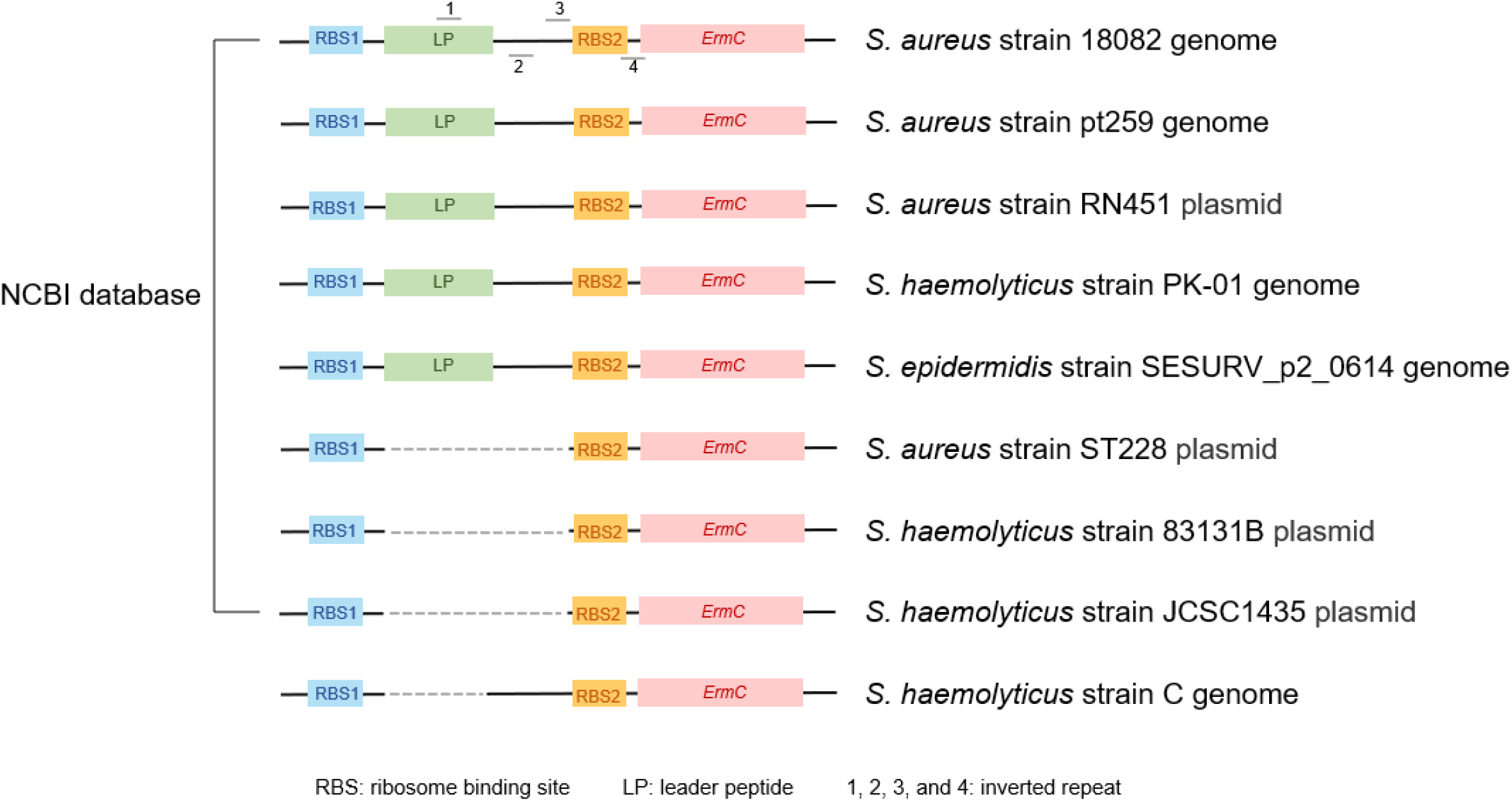
Identifying the target gene *ErmC*. Comparing the sequence of *ErmC* from strain C with its homolog in NCBI, and found a short leader peptide was missing from the upstream of *ErmC*.

**FIG 3.**
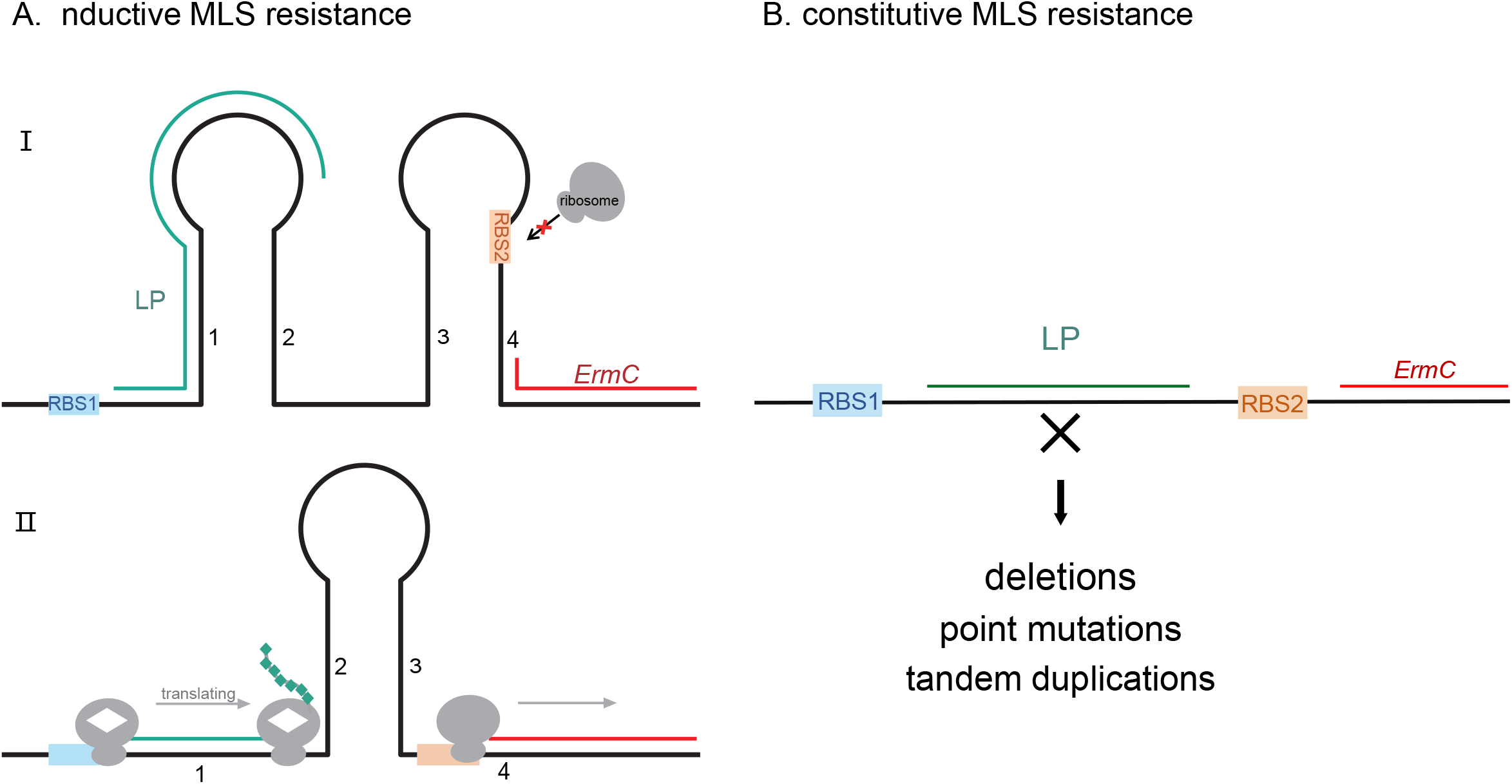
Mechanism of inducible (A) or constitutive (B) MLS resistance. (A) In situation I, *ErmC* mRNA is in an inactive conformation due to the structure of its 5’ end, which has a set of four inverted repeats paired as hairpin structure 1:2 and 3:4 that sequester the initiation sequences (ribosome binding site and initiation codon) for the methylase. Thus, the methylase cannot be synthesized, since the initiation motifs for translation are not accessible to ribosomes. In the presence of macrolide antibiotics (situation II), with the binding of erythromycin (white diamond) to a ribosome translating the leader peptide upstream of *ErmC*, the association between inverted repeat 1 and 2 is prevented. This favors the association between inverted repeat 2 and 3, which uncovers the initiation sequences to increase the translation efficiency of *ErmC*. (B) The variations of leader peptide decrease the stability of hairpin structure and render *ErmC* available for translation, thus result in constitutive MLS resistance.

### 3.3 Deletion of leader peptide upstream of *ErmC* resulted in MLS constitutive resistance of *S. haemolyticus*

To validate this hypothesis, we amplified the *ErmC* gene from *S. haemolyticus* strain C and constructed the recombinant plasmid pBT2-*ErmC*. A synthesized upstream leader peptide (short for LP) was inserted into the upstream of *ErmC* in plasmid pBT2-*ErmC* to form pBT2-LP-*ErmC*. The construction of recombinant plasmids was shown in Fig 4. Due to the absence of a widely used standard strain of *S. haemolyticus* for drug sensitivity test, we chose *S. aureus* as the experimental subject. *S. aureus* ATCC25923, which was sensitive to almost all antibiotics, usually used as a negative control strain for antimicrobial susceptibility test [31]. Two recombinant plasmids, pBT2-*ErmC* and pBT2-LP-*ErmC*, and the backbone plasmid pBT2 were transformed into *S. aureus* ATCC25923.

**FIG 4.**
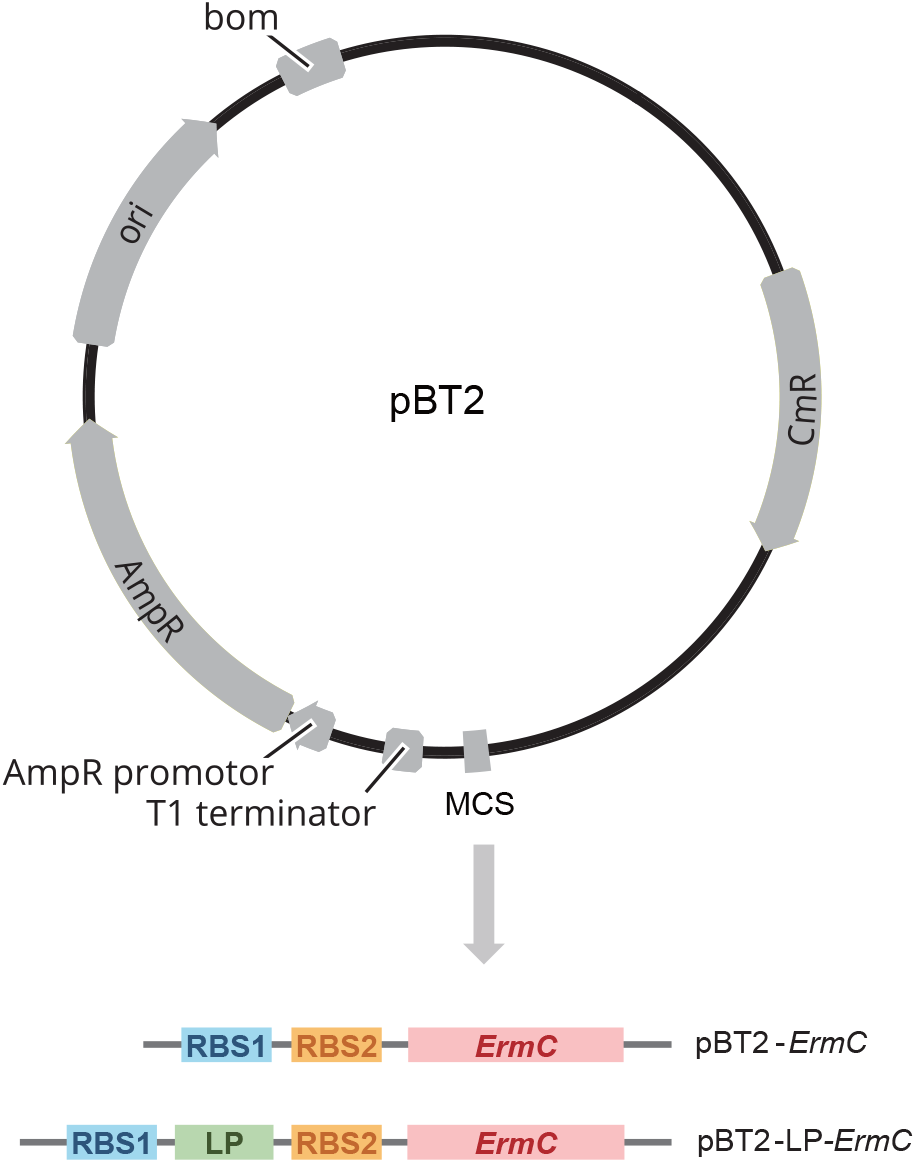
Schematic overview of plasmids construction. Plasmid pBT2-*ErmC* was constructed by digestion-ligation method with *ErmC* inserted into MCS of original plasmid pBT2. Plasmid pBT2-LP-*ErmC* was constructed by In-Fusion Cloning method, and the synthesized leader peptide was inserted into the upstream of *ErmC* in the pBT2-*ErmC*.

To evaluate the expression of *ErmC*, we extracted the total RNA of wild-type *S. aureus* and experimental groups transformed with different plasmids, followed by RT-PCR. Compared with wild-type *S. aureus* and *S. aureus* carried pBT2, the transgenic *ErmC* and LP-*ErmC* could express normally (Fig S3).

We performed a double disk diffusion test to determine MLS inducible/constitutive resistance, and the result is shown in Fig 5. *S. aureus* carried pBT2-*ErmC* was utterly unaffected by erythromycin or clindamycin, forming a phenotype of constitutive MLS resistance. Whereas the one harboring pBT2-LP-*ErmC* showed clindamycin resistance only on the side close to erythromycin susceptibility test disc (inhibition zone radius ∼7 mm), while the side far from the erythromycin disc remained clindamycin sensitive (inhibition zone radius ≈ 15 mm), resulting in the formation of a D-Circle, indicating that erythromycin induction was required for the resistance to clindamycin. Experimental validations revealed that the constitutive MLS resistance of *S. haemolyticus* strain C was due to the leader peptide deletion upstream of *ErmC*.

**FIG 5.**
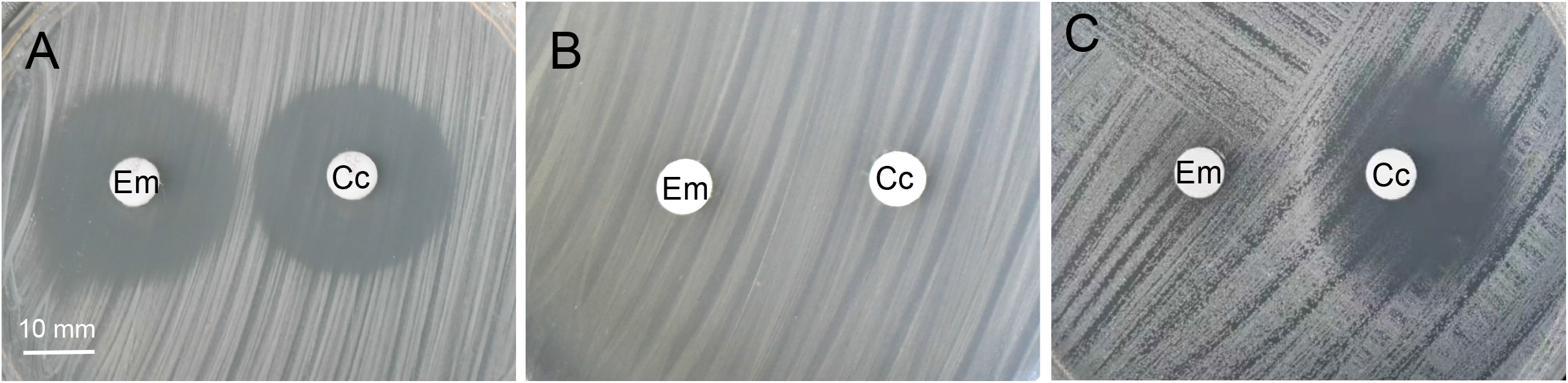
Results of antimicrobial susceptibility test of *S. aureus* ATCC25923. Em, erythromycin; Cc, clindamycin. (A) *S. aureus* carried backbone plasmid pBT2 were sensitive to both erythromycin and clindamycin. (B) *S. aureus* with pBT2-*ErmC* showed the constitutive MLS resistance. (C) *S. aureus* with pBT2-LP-*ErmC* showed clindamycin resistance only on the side close to erythromycin susceptibility test disc, thus resulted in the inducible MLS resistance.

### 3.4 The 23S rRNA methylation modified by *ErmC* may have adverse effects on bacterial growth

Multiple studies have shown that there were more constitutive MLS resistance phenotypes in the drug susceptibility testing of common pathogenic strains *Staphylococcus* and *Streptococcus* [32-35]. Since constitutive resistance mechanisms could be resistant to antibiotics substantially, we were wondering why bacteria evolved such inducible mechanisms. An interesting observation during our research may be the explanation. After the plasmids (pBT2, pBT2-*ErmC*, pBT2-LP-*ErmC*) were transformed into *S. aureus*, we found the growth rates varied considerably under the same culture condition. To confirm that, we determined the growth curve shown in Fig 6. The results showed that the growth of *S. aureus* carried pBT2-*ErmC* was significantly slower than others. We speculated that this observation might be due to the adverse effect of 23S rRNA methylation modified by *ErmC*. Compared with unmethylated ribosomes, the translation speed of methylated ribosomes was reduced, and protein synthesis was slowed down, directly affecting bacteria’s survival and metabolism, which impose a large fitness cost in the absence of antibiotics. Bacteria with constitutive expression of resistance genes can be against to bacteriostasis in face of antibiotic selection pressure. Whereas when the antibiotic pressure is relieved, the bacteria still express these nonessential genes may greatly aggravate the extra burden of strains. Therefore, bacteria have to possess the inducible resistance mechanisms for thriving in natural or few antibiotics environments.

**FIG 6.**
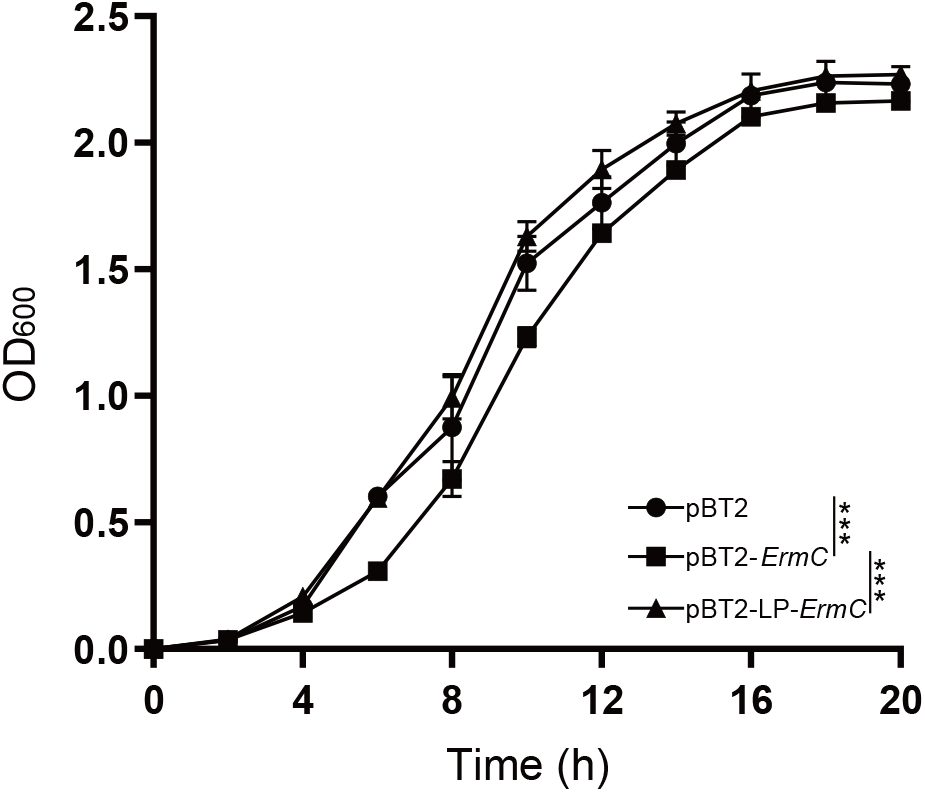
The growth curves of *S. aureus* ATCC25923 transformed with different plasmids. ANOVA was used for statistical analyses, *P < 0.05, **P < 0.01, ***P < 0.001, ****P < 0.0001.

## 4. Discussions

In this study, we acquired four *S. haemolyticus* strains isolated from clinical patients. While three isolates demonstrated antibiotic resistance as reported in previous studies (two were resistant to clindamycin; one was resistant to erythromycin), the other strain C manifested constitutive MLS resistance attracted our attention. Although such performance was reported in some other *Staphylococcus, Streptococcus* and *Enterococcus* [24-28], it was novel in *S. haemolyticus*. To confirm this discovery and illustrate the mechanism behind it, we acquired whole genome sequences of these four strains by NGS. After analyzing all potential antibiotic resistance genes, we found strain C contained a methyltransferase *ErmC* with a deletion of the upstream leader peptide, which may result in the constitutive MLS resistance ultimately, and this study was the first to identify such type of variation on *ErmC* gene in *S. haemolyticus*. The plasmids that mimic the wild-type and variated genotypes were constructed and transformed into *an S. aureus* antimicrobial susceptibility test strain. Based on the mRNA expression and drug resistance performance, it was clear that this small leader peptide was critical in regulating MLS resistance. However, further experiments can be extended to consummate this discovery, such as evaluating the protein level of *ErmC*, or identifying the dynamic changes in RNA secondary structure during the process.

As reported above, the constitutive MLS resistance of clinical pathogenic bacteria has become increasingly important, and the selection pressure of antibiotics mainly causes this situation. There are two main strategies to solve this crisis: limiting antibiotics or developing novel antimicrobials drugs. Taking advantage of the principle of fitness cost, restricting antibiotic usage can benefit the elimination of constitutive drug-resistant bacteria, because such bacteria will express some nonessential genes that may greatly aggravate the extra burden of strains and will be difficult to compete with others when antibiotic selection pressure disappears. In this case, the constitutive resistance bacteria will be quickly eliminated from the population, and the bacteria that remained from the competing challenges can be killed more easily and effectively.

It is widely known that *S. haemolyticus* is one of the most frequent pathogenic factors of staphylococcal infections [9]. Although it seems to lack the vital virulence elements compared with *S. aureus, S. haemolyticus* has the ability to acquire multi-resistance against various antimicrobial agents. It is noted that 75% of analyzed *S. haemolyticus* clinical isolates displayed multi-resistance [36]. Interspecies transfer of staphylococcal cassette chromosome mec (SCCmec) indicates that *S. haemolyticus* might be the reservoir of resistance genes [37, 38]. The genome plasticity of *S. haemolyticus* is characterized by many insertion sequences and SNPs, which may contribute to its acquisition of antibiotic resistance [39]. Here, the mechanism of antibiotic resistance revealed in this study may be another strong evidence to support that *S. haemolyticus* acts as a drug-resistance reservoir for other *Staphylococcus* and contributes to the emergence of epidemic clones of a more virulent nosocomial pathogen, *S. aureus*. In brief, multi-resistant strains with frequent intraspecific or interspecies gene transfer shorten the service life of antibiotics, more future studies can lead to our further understanding of the mechanisms in resistance genes’ transfer.

## 5. Conclusions

In summary, our study discovered a novel phenotype of macrolides/lincosamides resistance in *Staphylococcus haemolyticus* and clarified the mechanism behind it. Besides, we discovered that the expression of 23S rRNA methylase *ErmC* might inhibit the growth and metabolism of *S. haemolyticus*, suggesting to be the cost with no MLS pressure.

