## Supplemental Information for "Molecular basis and evolutionary cost of a novel phenotype of macrolides/lincosamides resistance in *Staphylococcus haemolyticus*"

**Supplementary Information**

**Table S1.** All potential resistance genes identified in *S. haemolyticus* strains ABCD.

**Table S2.** Summary of the reports about *ErmC* in NCBI database.

**Table S3.** Primers used in this study.

**FIG S1.** Schematic diagram of merge contigs in genome assembly process. The red peak represented coverage depth, the pure red or light blue line represented high or low depth respectively. Assemblies of five tools (SuperReads, BCALM, Tadpole, SPAdes, Megahit) were screened according to the coverage depth of reads, and those that were too high or too low were screened out. The trusted sequences with medium coverage depth were retained and merged.

**FIG S2.** Expression of *ErmC* in *S. aureus* ATCC25923. Five lanes represented marker, wide type, and strain carried plasmid pBT2, pBT2-*ErmC*, pBT2-LP-*ErmC*, respectively. The upper showed the target fragment (amplified from *ErmC*) and the lower showed the fragment of 16S rRNA internal reference. The length of target or internal reference fragment was 160 bp.

#### Supplementary Table 1

| Gene | *S. haemolyticus* Strains | | | | Gene Description | Antibiotic Resistance |
|---|---|---|---|---|---|---|
| | A | B | C | D | | |
| *AAC(6')-Ie-APH(2'')-Ia* | | | ■ | ■ | aminoglycoside N-acetyltransferase AAC(6')-Ie/ aminoglycoside O-phosphotransferase APH(2'')-Ia | aminoglycoside |
| *APH(3')-IIIa* | | ■ | ■ | | aminoglycoside phosphotransferase | aminoglycoside |
| *catA8* | | | ■ | | chloramphenicol acetyltransferase | phenicol |
| *dfrG* | | | ■ | | trimethoprim resistant dihydrofolate reductase dfr | diaminopyrimidine |
| *ErmC* | | | ■ | | Erm 23S ribosomal RNA methyltransferase | macrolide, lincosamide, streptogramin |
| *mecA* | ■ | ■ | ■ | | beta-lactam-resistant peptidoglycan transpeptidase | penam |
| *mphC* | | | ■ | ■ | macrolide phosphotransferase | macrolide |
| *msrA* | | | ■ | ■ | ATP-binding cassette (ABC) transporter | macrolide, lincosamide, streptogramin, tetracycline, oxazolidinone, phenicol, pleuromutilin |
| *PC1 beta-lactamase* | ■ | ■ | ■ | | blaZ beta-lactamase | penam |
| *SAT-4* | | ■ | ■ | | streptothricin acetyltransferase | nucleoside |
| *tet(45)* | | ■ | | | tetracycline efflux MFS transporter | tetracycline |
| *tet(K)* | | | | ■ | tetracycline efflux MFS transporter | tetracycline |
| *Vga (A)*$_{LC}$ | ■ | ■ | | | ATP-binding cassette (ABC) transporter | macrolide, lincosamide, streptogramin, tetracycline, oxazolidinone, phenicol, pleuromutilin |

#### Supplementary Table 2

| Species | *ErmC* with abnormal leader peptide | | *ErmC* with intact leader peptide | |
|---|---|---|---|---|
| | Chromosome | Plasmid | Chromosome | Plasmid |
| *S. aureus* | 16 | 42 | 7 | 63 |
| *S. epidermidis* | 1 | 14 | 2 | 4 |
| *S. haemolyticus* | 0 | 4 | 1 | 0 |
| Other *Staphylococcus* | 2 | 10 | 1 | 20 |
| *Streptococcus* | 1 | 2 | 4 | 10 |
| *Enterococcus* | 2 | 2 | 17 | 1 |
| Other bacteria | 9 | 9 | 4 | 1 |

#### Supplementary Table 3

| Primer Name | Primer Data |
|---|---|
| *ErmC*-XbaI-F | GCTCTAGAGCAGTATAAATTTAACGATCAC |
| *ErmC*-EcoRI-R | CGGAATTCCGATTCACAAAAAATAGGTACACG |
| In-fusion vector-F | GGTTATAATGAATCGTTAATAAGC |
| In-fusion vector-R | CTCTAGAGTCGACCTGCAG |
| In-fusion insert-F | CAGGTCGACTCTAGAGCAGTATAAATTTAACGATCACTCA |
| In-fusion insert-R | CGATTCATTATAACCACTTATTTTTTGTTTGGTTGATAAT |
| rt-*ErmC*-F | CACAGTCAAAACTTTATTAC |
| rt-*ErmC*-R | GGTCTATTTCAATGGCAGTTACG |
| rt-*Staphylococcus* 16S-F | CCATTGTAGCACGTGTGTAG |
| rt-*Staphylococcus* 16S-R | GAGATGTTGGGTTAAGTCCC |

### Supplementary Figure 1

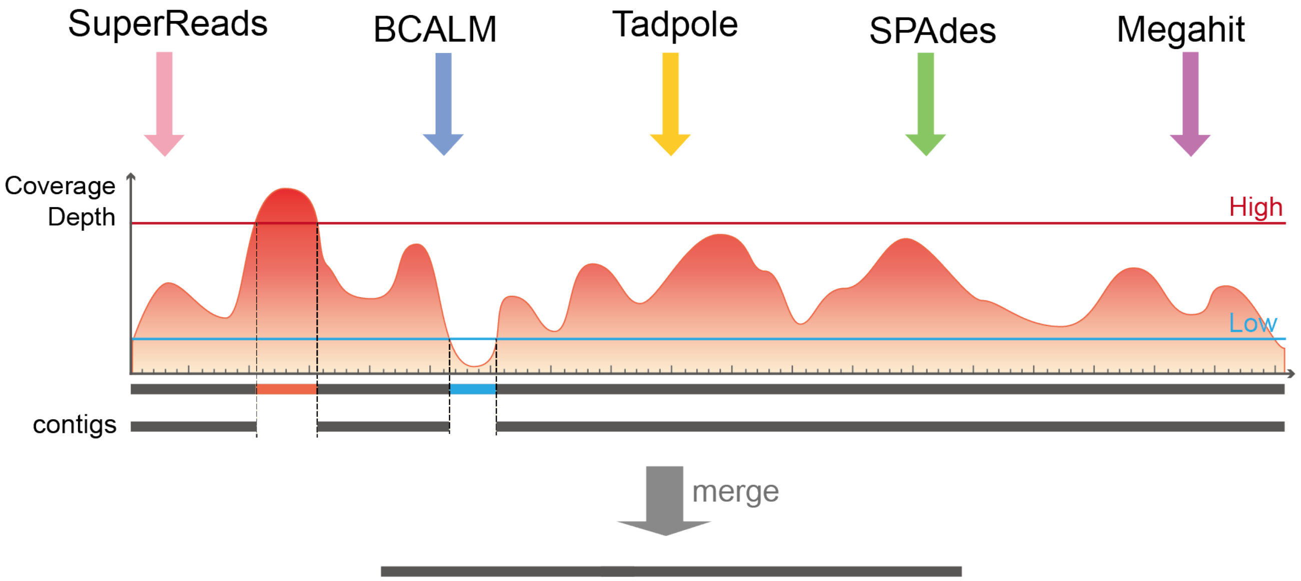

### Supplementary Figure 2

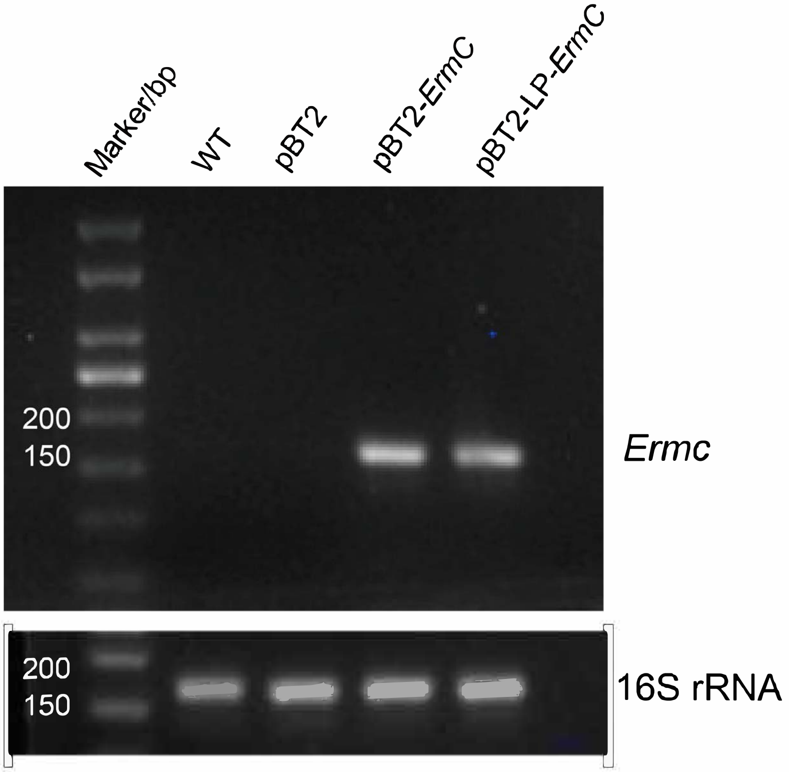
